# Long-term in vivo Monitoring of Gliotic Sheathing of Ultrathin Entropic Coated Brain Microprobes with Fiber-based Optical Coherence Tomography

**DOI:** 10.1101/2020.02.26.966887

**Authors:** Ian Dryg, Yijing Xie, Michael Bergmann, Gerald Urban, William Shain, Ulrich G. Hofmann

## Abstract

Microfabricated neuroprosthetic devices have made possible important observations on neuron activity; however, long-term high-fidelity recording performance of these devices has yet to be realized. Tissue-device interactions appear to be a primary source of lost recording performance. The current state of the art for visualizing the tissue response surrounding brain implants in animals is Immunohistochemistry + Confocal Microscopy, which is mainly performed after sacrificing the animal. Monitoring the tissue response as it develops could reveal important features of the response which may inform improvements in electrode design. Optical Coherence Tomography (OCT), an imaging technique commonly used in ophthalmology, has already been adapted for imaging of brain tissue. Here, we use OCT to achieve real-time, *in vivo* monitoring of the tissue response surrounding chronically implanted neural devices. The employed tissue-response-provoking implants are coated with a plasma-deposited nanofilms, which have been demonstrated as a biocompatible and anti-inflammatory interface for indwelling devices. We evaluate the method by comparing the OCT results to traditional histology qualitatively and quantitatively. The differences in OCT signal across the implantation period between the plasma group and the control reveal that the Parylene-type coating of otherwise rigid brain probes (glass and silicon) does not improve the glial encapsulation in the brain parenchyma.

## 1. Introduction

Implantable neural microelectrodes are developed to deliver therapeutic electrical stimulus for treating neural degenerative diseases, and to collect electrophysiology signals from surrounding neurons (Fanselow, Reid, & Nicolelis, 2000; Pinnell, Almajidy, Kirch, Cassel, & Hofmann, 2016; Pinnell, Pereira de Vasconcelos, Cassel, & Hofmann, 2018; Rouse et al., 2011; Tooker et al., 2014). These neural implants are designed to stay within the nervous system over a long period up to years (Jorfi, Skousen, Weder, & Capadona, 2015; Krüger, 2010; Nolta, Christensen, Crane, Skousen, & Tresco, 2015; Prasad et al., 2012). However, the performance of the electrode in terms of signal transferring efficacy and measurement signal-to-noise ratio deteriorate over time due to the inevitable foreign body reaction which eventually forms a dense glial sheath to encapsulate the implant, isolating it from the neural environment (Barrese et al., 2013; Kim, Hitchcock, Bridge, & Tresco, 2004; Polikov, Tresco, & Reichert, 2005; Szarowski et al., 2003; Ward, Rajdev, Ellison, & Irazoqui, 2009; Williams, Hippensteel, Dilgen, Shain, & Kipke, 2007).

To realize a long-term reliable neuro-electrode interface, it seems essential to minimize the foreign body response (Nguyen et al., 2014; Potter et al., 2014; Sohal, Clowry, Jackson, O’Neill, & Baker, 2016; Winslow & Tresco, 2010). To this end, many approaches have been proposed and investigated including innovations on bio-compatible flexible materials (Luan et al., 2017; Nguyen et al., 2014; Richter et al., 2013; Rousche et al., 2001; Rubehn, Lewis, Fries, & Stieglitz, 2010; Sohal et al., 2016; Stieglitz & Meyer, 1998; Takeuchi, Suzuki, Mabuchi, & Fujita, 2004; Böhler et al., 2017), decreasing implant size (Khilwani et al., 2017; Patel et al., 2015, 2016; Ferro et al., 2018) or surface modifications by coating the device with anti-inflammatory drugs that will actively release (Azemi, Lagenaur, & Cui, 2011; Boehler et al., 2017; Eles et al., 2017; Kolarcik et al., 2012; Potter et al., 2014).

These approaches are often evaluated by analyzing cellular changes corresponding to the progression of the foreign body reaction over time following implantation, based on postmortem immunohistochemistry (IHC) (Biran, Martin, & Tresco, 2007; Lo et al., 2018; Szarowski et al., 2003; Turner, 1999; Ward et al., 2009). The process of the foreign body reaction incorporates a rapid activation of microglial cells immediately following the implantation; and as the response develops up to three months, it reaches a chronic phase with compact astrocytic scar tissue generated to sheathe the implant (Leach, 2010, Pflüger et al., 2019). Normally, a large number of implanted and control animals are sacrificed to obtain sufficient temporal resolution of the progression curve as well as to reach a statistical significance. Furthermore, it is impractical to monitor the complete course of progression of the same object using postmortem histology.

*In vivo* two-photon microscopy (TPM) has been adopted for continuous monitoring the progression of the biological processes in neural tissue in response to foreign implants (Eles et al., 2017; Kozai, Eles, Vazquez, & Cui, 2016; Wellman & Kozai, 2018). It has been demonstrated that TPM is able to obtain spatiotemporal information of the dynamic cellular response to the implanted electrodes up to 12 weeks (Kozai et al., 2016). However, the imaging depth of TPM is limited within hundreds of micrometers (e.g. 200 μm), which only allows investigation of immune reactions very close to the brain’s surface.

Here we report our approach using fiber-based optical coherence tomography to monitor the development of the glial scar *in-vivo* over the course of the implantation. We previously demonstrated the use of fiber-based OCT to gain insight as a minimally-invasive imaging modality during in-vivo whole rat brain trajectories (Xie et al., 2013) and flexible implants (Xie et al., 2014), and elucidated the bases of its contrast’s origins (Xie et al., 2017). In this study, we first sought to compare the ability of fiber-based OCT with traditional IHC in observing the glial scar around another rigid implanted device (i.e. a silicon probe). Second, we used fiber-based OCT to investigate the foreign-body response of surface modifications by plasma deposited methane coatings, which have shown promise for affecting protein adsorption, reducing cellular adhesion, and reducing tissue encapsulation intravascularly in pigs (M. Bergmann, Ledernez, & Urban, 2015a; Michael Bergmann, 2015).

## 2. Materials & Methods

Eight female Sprague Dawley rats were each implanted with one inactive silicon probe and one OCT probe at ninety degrees to one another as shown in Figure 1. To assess glial scar development around silicon probes, OCT monitoring was performed weekly for 8 weeks before sacrificing for histological assessment with IHC. To investigate the effect of plasma deposited CH_4_ coatings on glial scar development around OCT probes, OCT probes were either CH_4_-coated or uncoated. Then, tissue properties were assessed by measuring the tissue attenuation coefficient of the OCT signal (Xie et al., 2013, 2014) and finally by IHC.

**Figure 1:**
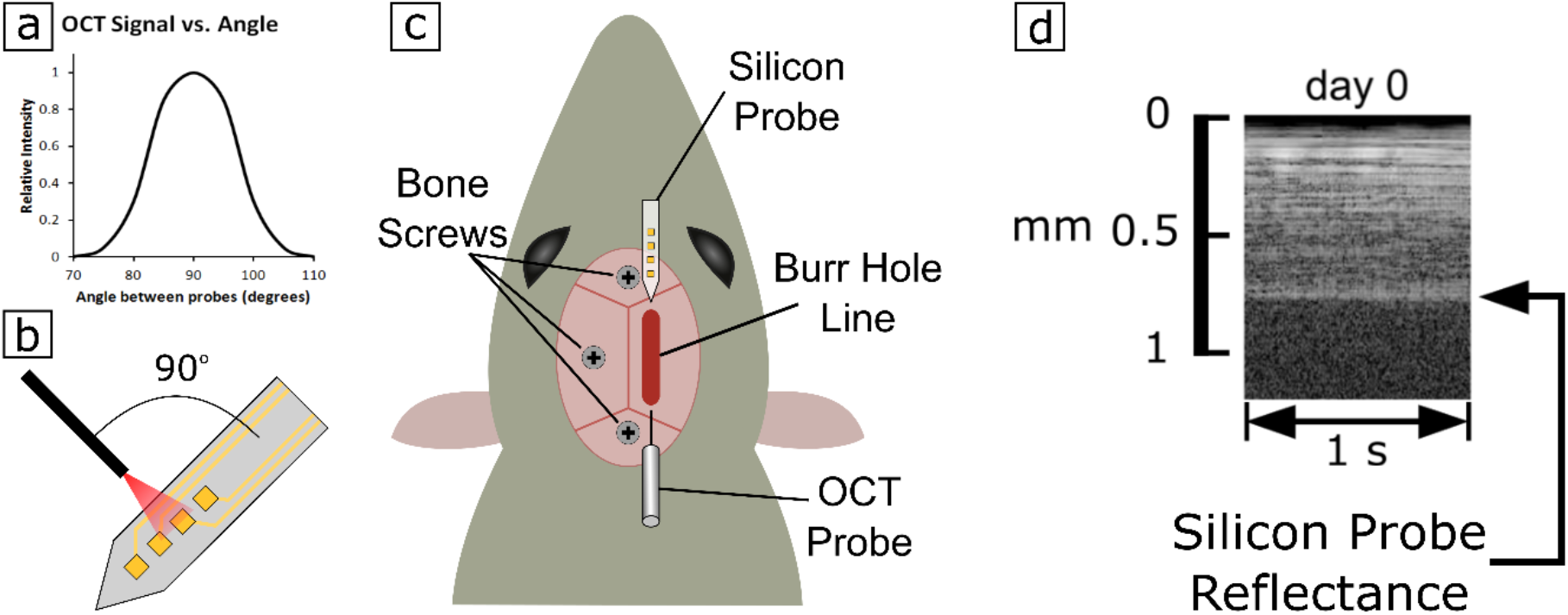
Surgical setup. (a,b) To detect the silicon probe with the OCT signal, probes had to be angled as close to 90 degrees from one another as possible, maximizing reflectance back into the OCT probe. (c) Top view of the surgical setup. A burr hole line was drilled to allow implantation of probes into brain tissue. The OCT probe and silicon probe were pre-aligned before advancing into brain tissue. The silicon probe was advanced first, followed by the OCT probe until the silicon probe could be detected in the OCT signal. (d) OCT signal during surgery after both probes are aligned and implanted, showing the reflectance of the implanted silicon probe.

### 2.1 Implantable probes: optical coherence tomography fiber probe and silicon microelectrode probe

The customized OCT fiber probe (SM800-CANNULA, Thorlabs) consists of a ceramic fiber connector ferrule (CF126-10, Thorlabs) with 2.5 mm diameter and 10.5 mm length and an 8 mm long single mode fiber (SMF) (ø 125 μm, SM800-5.6-125, Thorlabs) with clean cleaved distal end. The single mode fiber is implanted into brain tissue to enable monitoring foreign body reaction *in situ*, while the ferrule remains exposed and is connected with the SMF cable of the OCT system each time the OCT measurement is conducted. The silicon-based rigid microelectrode probe is chosen in this study as it has a wide range of applications in neuroscience. Thus, a spatiotemporal *in situ* assessment on the foreign body reaction against the chronically implanted silicon probe would be of great interest to neuroscientists. The silicon probe used here is 50 μm thick, 140 μm wide and 10 mm long. Both OCT fiber probes and silicon microelectrode probes are immersed in 70% ethanol for 20 min for disinfection, and rinsed thoroughly with sterile saline (0.9% Sodium Chloride) before implantation.

### 2.2 Plasma Deposition of CH_4_ Coating on Glass OCT Probes

Using methane as the starting material, silicon probes were plasma coated as previously described (M. Bergmann, Ledernez, & Urban, 2015b; Ledernez, 2011). Briefly, a magnetron enhanced plasma polymerization process was conducted in a system from the company Shinko Seiki, with two parallel titanium electrodes separated by 10 cm. Samples were rotated between the electrodes to ensure good homogeneity of the resulting plasma films. The plasma chamber was evacuated down to a pressure of 0.1 Pa to avoid cross-contamination, then coated at a pressure of 5 Pa using a power of 45 W.

### 2.3 Implant Surgery

All experiments in this study were performed with approval from the locally responsible Animal Welfare Committee with the Regierungspräsidium Freiburg in accordance with the guidelines of the European Union Directive 2010/63/UE under permit G13/51. We used 8 adult female Sprague Dawley rats (Charles River, Germany) weighing 280-320 g (four rats in each group). One silicon probe and one OCT probe were implanted under micromanipulator control into each animal. To study the effect of CH_4_ plasma coating (M. Bergmann et al., 2015a; Michael Bergmann, 2015) on the foreign body reaction, four of the rats were implanted with CH_4_ plasma coated OCT probes. The remaining OCT probes and all of the silicon probes were uncoated.

The surgical procedure was performed as previously described (Richter et al., 2013). In brief, rats were anesthetized (induction: 5% isoflurane in 2L/min O_2_; maintenance: 0.5-3% isoflurane in 2L/min O_2_), the surgical area was shaved, and rats were placed on a water-circulating heating pad to maintain a body temperature of 35 °C. The rats were fixed into a stereotactic frame (Kopf Instruments), and eyes were covered with ophtalmic ointment. The surgical area was sterilized using alternating wipes of polyvidone-iodine and alcohol. A midline incision was made along the scalp, and the skin was pulled aside to expose the skull’s surface. Bregma was identified and used as a landmark coordinate (0,0mm). Bone screws were placed into the skull anterior to Bregma, posterior and lateral (left) to Bregma, and posterior to Lambda, to better secure a headcap with dental acrylic later. A burr hole line 9mm long and 1mm wide was drilled from 1mm posterior and 3mm lateral (right) to 10mm posterior and 3mm lateral (right) of Bregma. Then, using two different arms on the stereotactic frame, the implantable OCT imaging probe and the silicon probe were positioned above the burr hole line, each angled 45 degrees up from horizontal, and angled 90 degrees from one another (Fig. 1). The OCT probe was connected to the OCT imaging system, and the OCT probe and silicon probe were aligned laterally, pointing the OCT probe at the silicon probe ~1mm away, until the strongest reflectance signal from the silicon probe was achieved in the OCT signal. This process was essentially a practice run in air, creating the correct alignment and positioning of the probes for best imaging signal when implanted into tissue. Once correctly aligned in the medio-lateral direction, the probes were separated along the antero-posterior direction, and the silicon probe was advanced into brain tissue ~10mm, maintaining the same angle and lateral positioning as during the alignment in air. Then, the OCT probe was slowly advanced into brain tissue, until the OCT reflectance signal from the silicon probe was detected ~1-2mm away from the OCT probe tip. The OCT probe was advanced until the distance between the OCT probe tip and the silicon probe was 1mm, as measured by the OCT signal. The entire burr hole line including the entry points of the probes were covered in gelfoam (Pfizer) wetted with sterile saline, then with silicone elastomer (Kwik-Sil, World Precision Instruments), and finally sealed with dental acrylic. Dental acrylic was sequentially applied in several layers, securing the implanted probes in place, and building the dental acrylic headcap to cover the wound of the animal. The ferrule interfacing between the implanted OCT probe and the fiber optic cable leading to the OCT imaging system was left uncovered from dental acrylic, enabling access for connection to the OCT imaging system throughout the study. Carprofen (4 mg/kg) was administered subcutaneously for 5 days post-surgery to manage pain. The animals were housed separately with enrichment and daily inspection.

### 2.4 Optical Coherence Tomography

The OCT imaging system used in this study is identical to that which is previously described (Xie et al., 2013, 2014). The fiber-based spectral radar OCT system utilizes a superluminescent diode with center wavelength at 840 nm as light source. The system axial resolution is approximately 14.5 μm measured in air. Briefly, we used OCT (Fercher, Hitzenberger, Drexler, Kamp, & Sattmann, 1993) to monitor the development of foreign body capsule formation around implanted OCT fiber probes and silicon probes in brain tissue. The implanted fiber transmits incident light into brain tissue and collects light that scatters back into the fiber. The intrinsic optical properties of the tissue are encoded by the interference pattern created by the incident and backscattered light, which is detected by a spectrometer to construct A-scan (1-dimensional depth scan) signals that present backscattered light intensity as a function of depth. As our OCT utilizes a low-coherence superluminescent light source in the near infrared (840 nm) contrast depends on the attenuation (optical density of tissue and cells) and the dichroitic properties (like myelin fibers) of the illuminated tissue (Kut et al., 2015). Both sending and receiving optics are defined by a cleaved single mode fiber (0.22 NA, SM800-5.6-125, Thorlabs) achieving extremely high localisation of beams and thus making perfect arrangement an important requirement.

Implanted OCT fibers were either coated with plasma-deposited CH_4_ or left uncoated. After a CH_4_-coated or uncoated fiber had been implanted and fixed in place within the dental cement headcap, the fiber optic cable connecting the implanted fiber to the OCT imaging system could be disconnected, allowing the animal to move around freely. After disconnecting the implanted OCT probe, the ferrule was wrapped in parafilm for protection. Before each imaging session, the parafilm was removed and the ferrule was cleaned using Fiber Connector Cleaning Solution (FCS3, Thorlabs). The implanted OCT probe was then connected to the OCT system via the fiber optic cable and *in vivo* imaging was performed. If a poor signal was observed, the fiber optic cable was removed, the ferrule re-cleaned, and the OCT system re-connected to try again. Rats were not anesthetized during OCT imaging, but only data recorded while the animal was motionless were used for analysis, as movement of the animal can disrupt the OCT signal quality. To create an OCT signal profile, imaging data over an entire second of motionless collection was averaged together and plotted as OCT signal intensity vs. distance from the tip of the OCT probe.

### 2.5 Tissue Histology

Rats were given an intraperitoneal injection of ketamine/xylazine and sacrificed by cardiac perfusion with PBS to clear the blood, and then 4% paraformaldehyde to fix the tissue. Rat heads were removed and post-fixed in 4% paraformaldehyde overnight, then washed in PBS and stored in HBHS with sodium azide prior to brain removal. Because we wanted to compare the OCT signals to tissue histology, we aimed to maintain both the implanted OCT imaging probe and silicon probe *in situ* within one slice, as demonstrated previously (Woolley, Desai, & Otto, 2013; Woolley, Desai, Steckbeck, Patel, & Otto, 2011). To do this, headcaps were carefully drilled into using a surgical drill, and implants were cut at the entry point through the skull to maintain implant positioning within brain tissue. Tissue blocks were carefully aligned for sagittal slicing, keeping both probes along the plane of slicing. Using a vibratome, tissue blocks were supported with agar gel and sliced at 300μm thickness to capture both probes within a single slice. Slices were immunolabeled using primary antibodies GFAP (1:1000 dilution, rat IgG2a, Invitrogen cat# 13-0300) to label astrocytes, Iba1 (1:250 dilution, rabbit IgG, Wako cat# 016-20001) to label microglia, NeuroTrace 640/660 (1:250 dilution, Invitrogen cat# N21483) Nissl stain to label neuronal cell bodies, and Hoechst 33342 (1:1000 dilution, ThermoFisher) to label cell nuclei. Secondary antibodies goat anti-rat AlexaFluor 488 (1:200) and goat anti-rabbit AlexaFluor 594 (1:200) were used to target GFAP and Iba1 primaries, respectively. Slices were mounted in Fluoromount G and cover slips were sealed with nail polish. Slides were imaged at 10X and 30X using an Olympus spinning disc fluorescence microscope, scanning across multiple imaging fields to image the entire area of interest. Optical sections were taken every 0.3μm throughout the thickness of the slice to gather fluorescence data from the whole tissue slice. Background subtraction was performed, and a custom MATLAB script stitched the scanned image stacks together to create a 3D representation of the fluorescence data. Finally, all optical sections were compiled together into one image by taking a maximum projection. All images were captured using the same settings across fluorescence channels.

To assess the effects of CH_4_ plasma coating on the foreign body reaction, the fluorescence of glial cell markers around implants were examined. Analysis of the glial cell responses surrounding implants were achieved using a custom MATLAB script that measured the average fluorescence intensity of each channel versus distance from the implant in 10X images. Fluorescence profiles from all animals in each group were averaged together, normalized to control fluorescence and subjected to statistical analysis (student’s t-test) compared to control tissue. Control data was gathered by averaging the fluorescence intensity of a field far (>2mm) from implants but at the same tissue depth. As an additional assessment of the effect of the CH_4_ plasma coating, the OCT signal’s attenuation factor was calculated for each probe.

To compare the OCT imaging signal to the IHC, 30X image scans of tissue between the OCT probe and the silicon probe were collected. In those images, the shape of the light collected and imaged by the OCT probe was recreated. The OCT probes were composed of only a single mode fiber (5.6 μm diameter core, numerical aperture = 0.12) with an acceptance angle of 6.9 degrees. Thus, the tissue imaged by the OCT probes was a conical shape starting at the OCT probe tip and diverging at an angle of 6.9 degrees in both directions. The IHC images were analyzed by measuring fluorescence profiles from the tip of the OCT probe outwards in a conical shape, rotating the angle of the profiles to cover the whole area of the cone, and then averaging those profiles together to create one profile. In this way, the OCT imaging signal could be compared to the analogous fluorescence profile from the same tissue that the OCT system imaged (Figure 2)

**Figure 2:**
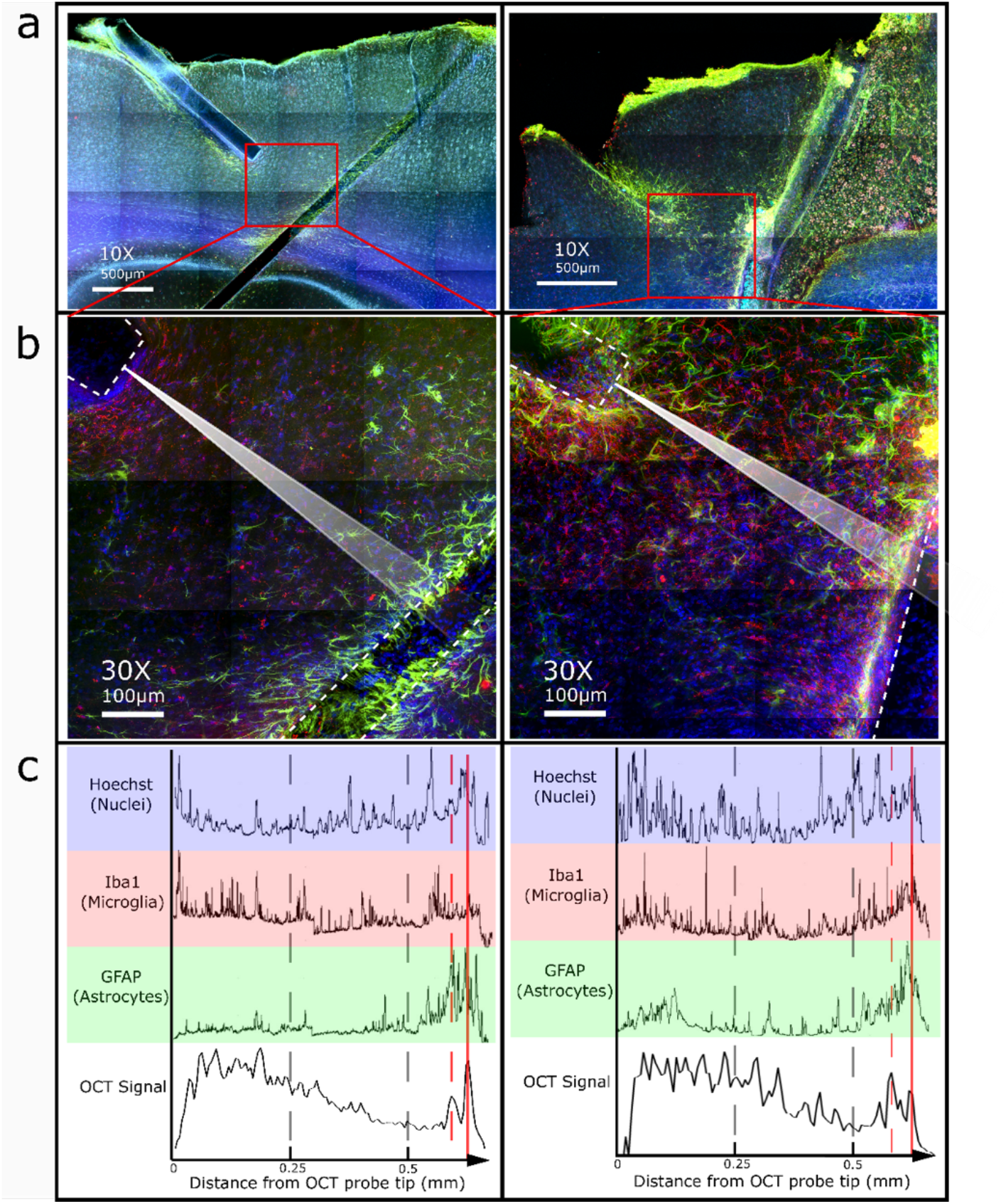
Comparisons between IHC and OCT signal. (a) 10X scans show implant positions of two representative animals (scalebar: 500μm). (b) 30X scans of the tissue that had been imaged by OCT (illustrated by white beam) allows comparison between OCT signal and IHC profiles. Implants are outlined by dotted lines (scale bar: 100μm). (c) OCT signal profile plotted below fluorescence profiles from the same tissue that had been imaged by OCT. The x-axis spans from the OCT imaging probe tip, through the tissue, and ends at the implanted silicon probe. Increases in fluorescence intensity profiles from labeled glial cells (dotted red line) match with the increase in OCT signal at the location of the silicon probe (solid red line).

## 3. Results and Discussions

### 3.1 Comparison of OCT Signal to Tissue Histology by IHC

When comparing the OCT signal to the fluorescence profiles from the tissue histology images, an increase in both OCT signal and fluorescence of glial cell markers were observed around the implanted silicon probe (Figure 2). However, no other correlations could be drawn between signals.

The location of the glial scar in fluorescent plots matched the location of the implanted silicon probe in the OCT signal. However, we were unable to find a difference in the extent of glial scarring around the silicon probes over time using the OCT signal. From (Xie et al 2017), it was determined that OCT image contrast in brain tissue is more likely from fibrous, well-ordered, myelin-rich structures such as axons, than from other cells such as glia. This helps to explain why we were unable to detect changes in gliosis over time using OCT signal.

Interestingly, we were able to detect changes in the silicon probe’s position relative to the OCT probe (Figure 3). This may be an important observation since electrode position relative to nearby neurons greatly affects the signals recorded by that electrode.

**Figure 3:**
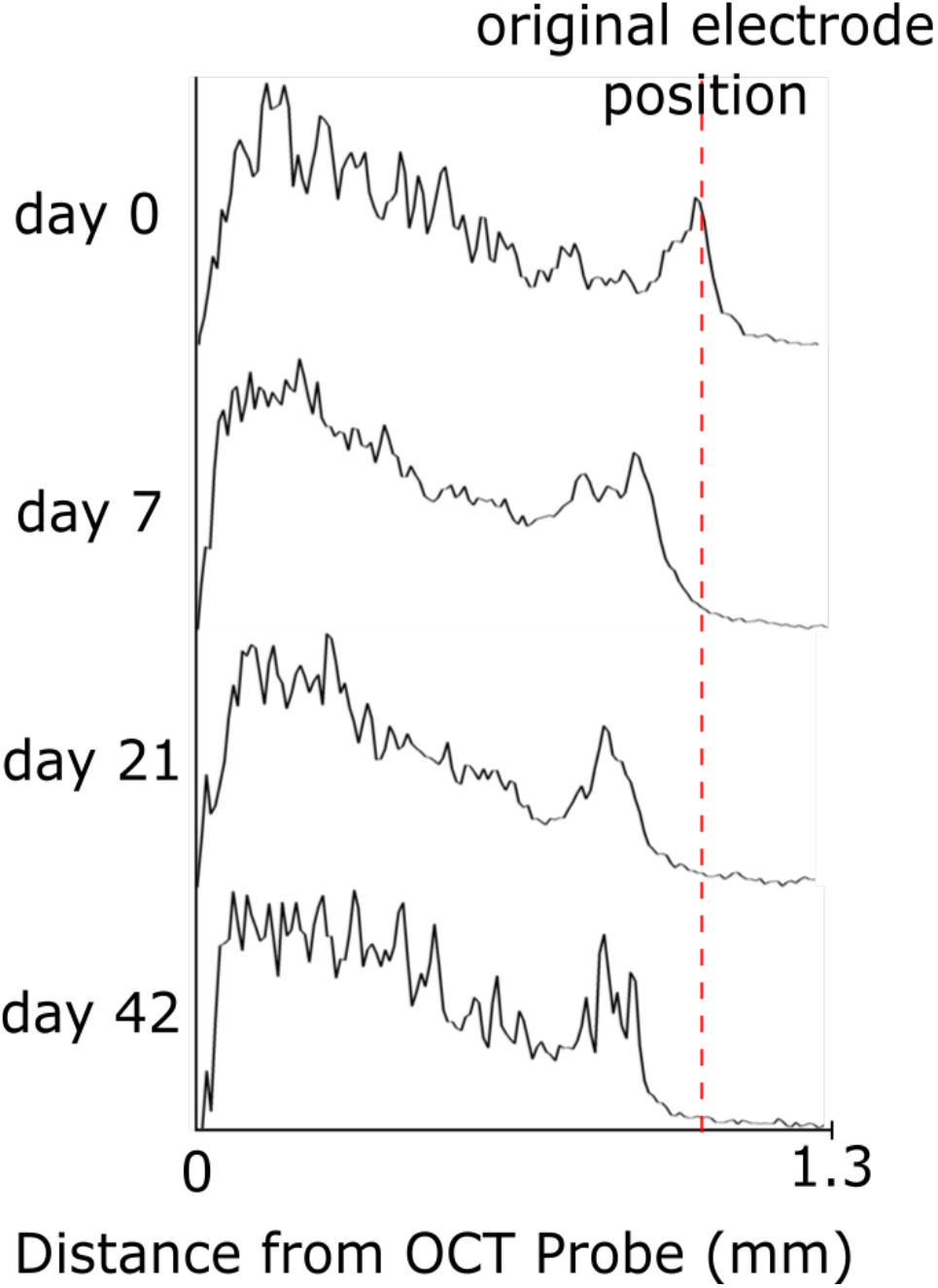
Change in electrode position can be detected and measured by the OCT signal profile.

### 3.2 Effect of CH_4_ Plasma Coating on Foreign Body Reaction

There were no trends or significant differences in tissue attenuation factor or maximum signal intensity from the OCT imaging data between the CH_4_-coated and uncoated groups (Figure S1, S2). Additionally, histological analysis by IHC revealed no significant differences between coating groups (Figure 4). Both coated and uncoated OCT probes showed significantly higher GFAP (Figure 4a) and Iba1 (Figure 4b) fluorescence around the implants compared to control tissue greater than 2 mm away from any implant. However, GFAP and Iba1 fluorescence were no longer significantly different from control tissue at a shorter distance from CH_4_-coated probes than for uncoated probes (Figure 4c).

**Figure 4:**
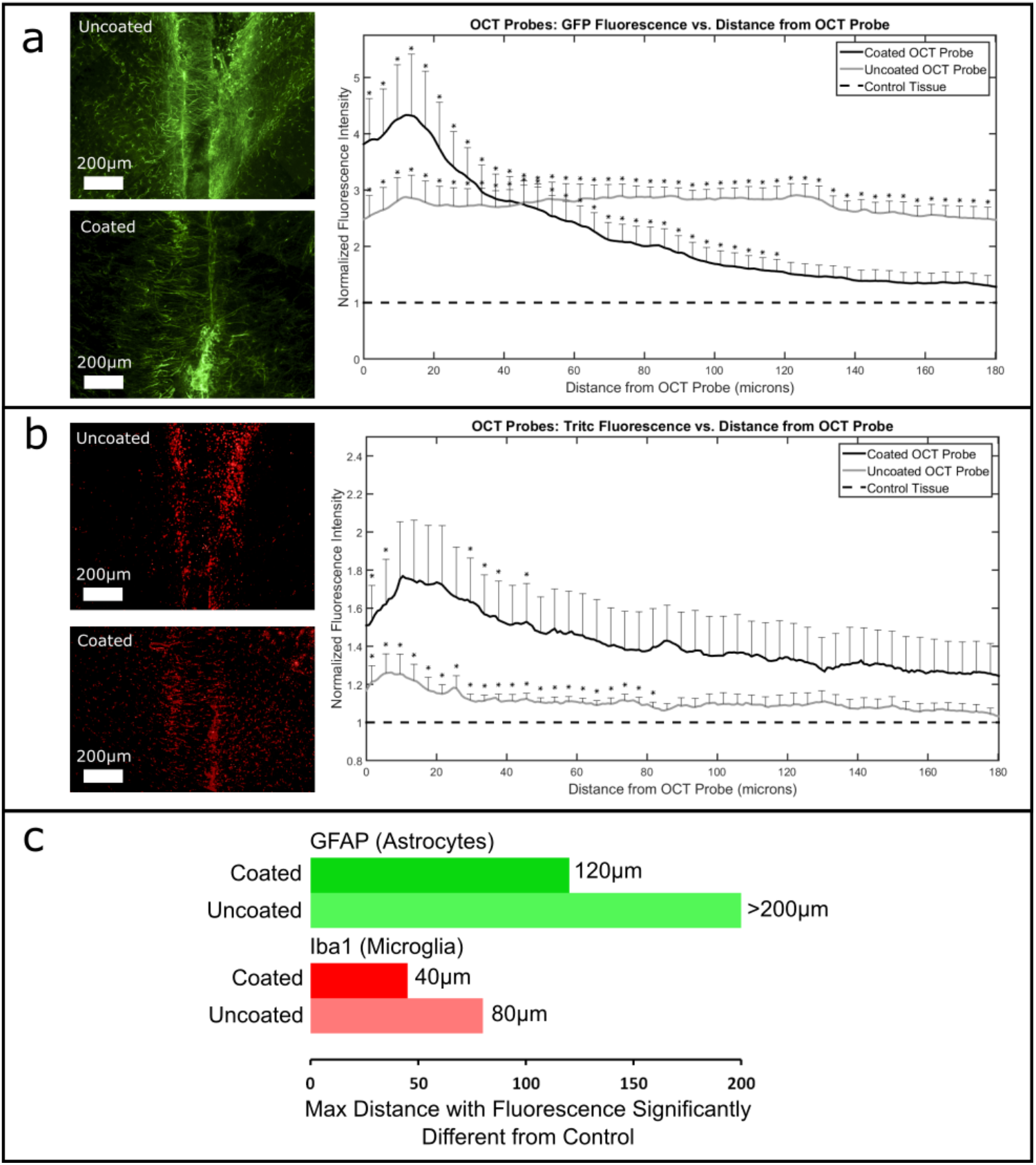
Effect of CH_4_ plasma coating on glial encapsulation of the OCT probes. Shown are quantified encapsulation by (a) astrocytes labeled by GFAP, and (b) microglia labeled by Iba1, with representative uncoated and coated OCT probe images. The maximum distance with fluorescence intensity significantly different from control tissue is plotted in (c).

In a previous implant study, CH_4_ plasma coatings reduced tissue encapsulation of titanium intravascular implants (Michael Bergmann, 2015). However, our study revealed no clear differences between CH_4_-coated or uncoated probes by OCT Imaging or by IHC. This may be explained by the differences between the intravascular and nervous environments, and by the flow component present intravascularly but not in brain tissue. Also, the aforementioned study used titanium as the base material, whereas this study used a glass fiber. The fluorescence of CH_4_ coated probes stopped being significantly different from controls closer than uncoated probes, suggesting the glial capsule is smaller around CH_4_ coated probes. However, the variance was generally high for both groups. Further study with larger sample sizes should be performed to elucidate any relationship between CH_4_ plasma coatings and the foreign body reaction in the brain.

## Acknowledgements

Part of this study was supported by the Cluster of Excellence Brainlinks-Braintools (EXC 1086).

---

**Figure S1:**
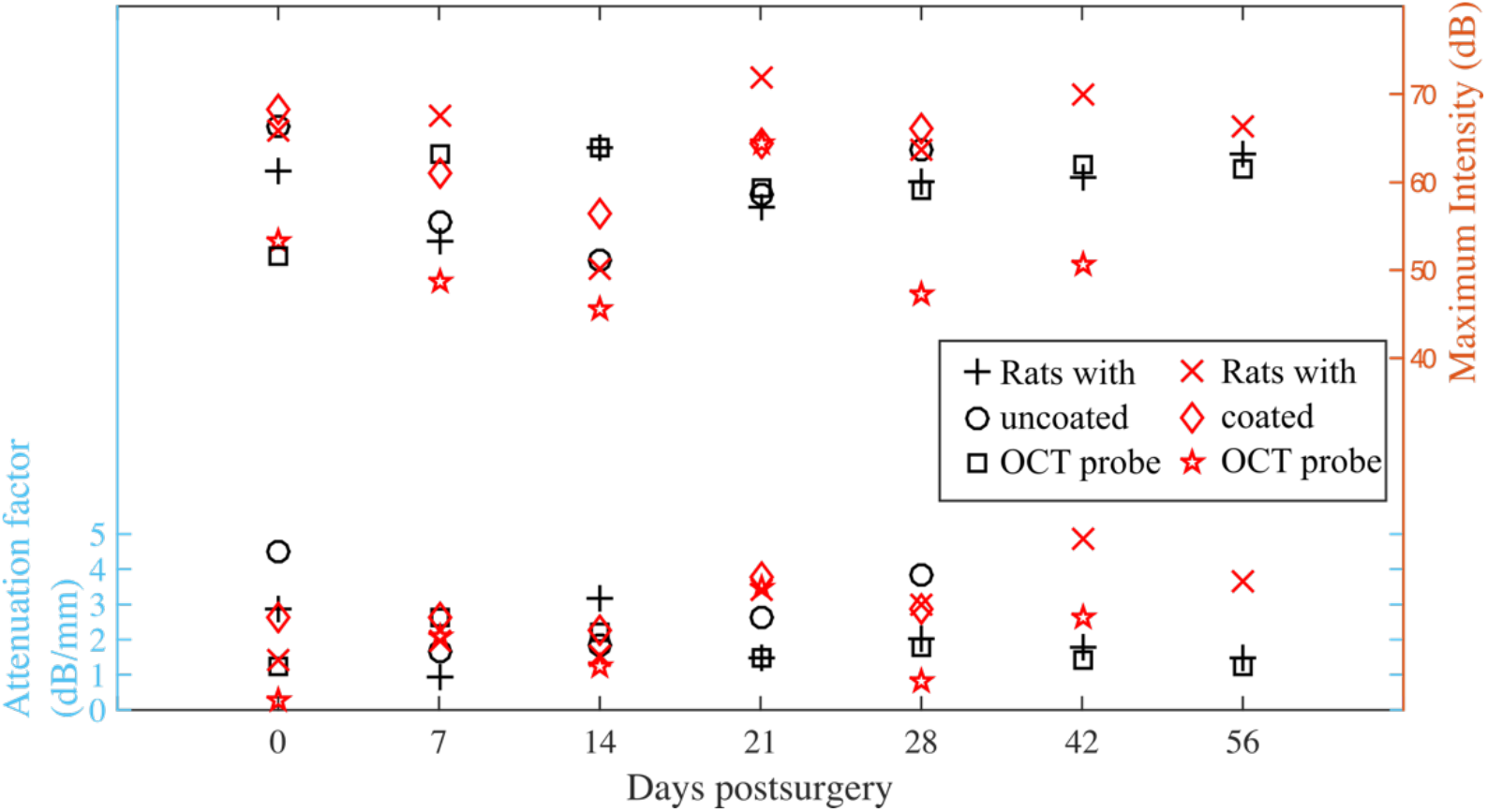
Attenutation factor and maximum intensity that are extracted from OCT measurements do not illustrate markable differences between rats with coated and uncoated OCT probes across the entire implantation period. There is a consistent trend between the two groups that is the gliotic sheathing process reaches its maximal at day 21 post implantation, as shown in a peak value of both attenuation factor and maximum intensity.

